# Alignment of Artificial Neural Networks with Human Eye Movement Patterns in Facial Expression Recognition

**DOI:** 10.64898/2025.12.12.693899

**Authors:** Meng Liu, Lihui Wang

**Author notes:** Corresponding author: Dr. Lihui Wang, School of Psychology, Shanghai Jiao Tong University, 1954 Huashan Road, Shanghai 200030, P. R. China.

## Abstract

Artificial neural networks (ANNs) have become powerful tools for modeling human perception and hehavior, yet it remains unclear if they resemble the sequential eye movement strategies humans employ during visual processing. Human studies demonstrate that the first two fixations play distinct roles in face processing: the initial fixation (Fix I) captures global configurational information, while the second (Fix II) targets local diagnostic details, with the two working collaboratively to support recognition. In this study, we tested if such properties emerge in ANNs trained for facial expression recognition. We fine-tuned a convolutional neural network (VGG16) and a Vision Transformer (ViT) on multiple facial expression datasets and evaluated their performance on the same images used in a human eye-tracking experiment. Human participants (N = 28) achieved a mean accuracy of 84.2%, VGG reached 82.9%, and ViT outperformed both at 90.0%. Perturbation analyses showed that both models aligned with the independent functions of Fix I and Fix II, but only ViT displayed a synergistic effect when information from both fixations was combined. Layer-wise analyses revealed stronger correspondence with Fix II than Fix I in both models, while ViT uniquely captured the statistical dependencies of fixation transitions. These findings suggest that VGG primarily reflects feedforward, detail-based processing, whereas ViT supports feedback-like integration across sequential fixations. Our results indicate that current ANNs can approximate human eye movement patterns in face processing, and highlight the potential of transformer-based architectures to model sequential strategies in human visual exploration.

## Introduction

Artificial neural networks (ANNs) have become powerful tools in understanding how human cognition works^1–4^. Training an ANN for a specific task and producing human-like behavior enables the modeling and hypothesis-testing of the cognitive processes involved in that task. Among the tasks where ANNs achieve or even exceed human-level performance, face processing has been extensively investigated in recent years, providing important insights into the mechanisms underlying this essential visual function in daily life and social communication^5–8^. However, to infer human cognitive mechanisms from human-like behavior in ANNs, a fundamental question arises: to what extend do humans and ANNs share the same processing pipeline?

One typical question in face processing is how facial information is optimally obtained and integrated to achieve efficient and flexible behavior. For instance, recognizing the identity of a friend in front of you may primarily rely on a distinctively high nose bride and pointy chin, whereas judging the emotional state of the same person may rely more on the eyes and mouth. One approach to modeling this information usage is to build a function between various facial features – such as action units in facial expression studies^9^ – and recognition performance. Comparing such stimulus-response functions between humans and ANNs can thus reveal the extent to which ANNs capture the algorithm of information processing in human face recognition^10,11^.

Despite receiving the same stimulus input and reproducing the response output, a fundamental bodily constraint on human visual perception is missing in ANNs – the sequential eye movements used to obtain visual information. Due to the biological architecture of the eyes, even in common daily task such as face processing, the required information cannot be fully acquired in a single fixation. This constraint, however, drives the optimization of fixation sequence into a parsimonious and collaborative sampling strategy developed through experience^12–14^. In a recent study^15^, we showed that the initial two fixations^16^ have distinct but collaborative functions in sampling facial information. Specifically, across both identity and emotion recognition tasks, the first fixation (Fix I) typically landed near the face center to capture global configurational information, while the second fixation (Fix II) diverged to key regions such as the eyes and mouth to analyze local details. Importantly, although the pattern of Fix II alone – but not Fix I alone – predicted recognition performance, the combined pattern led to better prediction, suggesting a functional collaboration between the two fixations^15^.

Both ANNs and Human fixation patterns are shaped by task optimization, raising the question of if ANNs exhibit properties similar to human fixations sequences in face recognition. Answering this question would not only test a core assumption of modeling human cognition with ANNs but also shed light on whether an artificial system lacking a human body can exhibit aspects of embodied cognition. Moreover, unlike information processing inferred from stimulus-response functions, eye movement patterns provide grounding for cognitive processes in at least two respects. First, fixation locations on the face image indicate the importance of specific information for the task and its associated brain functions. Notably, a patient with amygdala damage who failed to recognize fearful expressions was found to lack fixations on eye region^17^. When fixations were explicitly directed to the eyes, the ability to perceive fear was restored. Second, transitions between fixation locations reflect predictive processing of visual information^18,19^. Given the same overall fixation locations, an altered or non-preferred order can impair recognition performance^20,21^.

To address the above question, we investigated if patterns similar to human fixation sequences would emerge in ANNs trained for face processing. We adopted two widely used vision models: the Visual Geometry Group model(VGG)^22^, a popular deep convolutional neural network for face recognition tasks, and the Vision Transformer(ViT)^23^, a state-of-the-art model inspired by large language models^24^. We trained both ANNs to perform a facial expression recognition task used in a previous study^15^, reaching human-level prediction accuracy. Based on human fixation data in this task, we first examined whether and how perturbing fixation-related visual information in face images would affect the models’ performance. We predicted that, if the ANNs used a strategy similar to human fixation sequences, perturbing information covered by Fix II would have a greater effect than Fix I. Crucially, to test the collaboration between fixations, we asked if the perturbation effect based on the combined information from both fixations would exceed the sum of their individual effects – that is, a synergistic effect. We further predicted that this synergistic effect would be more pronounced in ViT due to its self-attention mechanism, which captures statistical dependencies in visual input. In a second step, we computed the density map for each fixation and tested its alignment with the activation maps derived from the model layers. We predicted stronger alignment with the density map of Fix II than Fix I. Finally, we quantified the transition probability between sequential fixations and tested if the transitional pattern aligned with information transitions captured by the attention mechanism in ViT.

## Results

### Emotion recognition in human participants and ANNs

In the human experiment, 28 participants were presented with front-view face images during which their eye movements were recorded. Each image displayed a facial expression from one of the seven categories: Angry, Disgust, Fear, Happy, Neutral, Sad, or Surprise. Participants were asked to judge the emotion by selecting the correct label from the seven alternatives. The mean recognition accuracy across all participants was 84.2%. The confusion matrix for emotion recognition is shown in Figure 1A.

**Figure 1.**
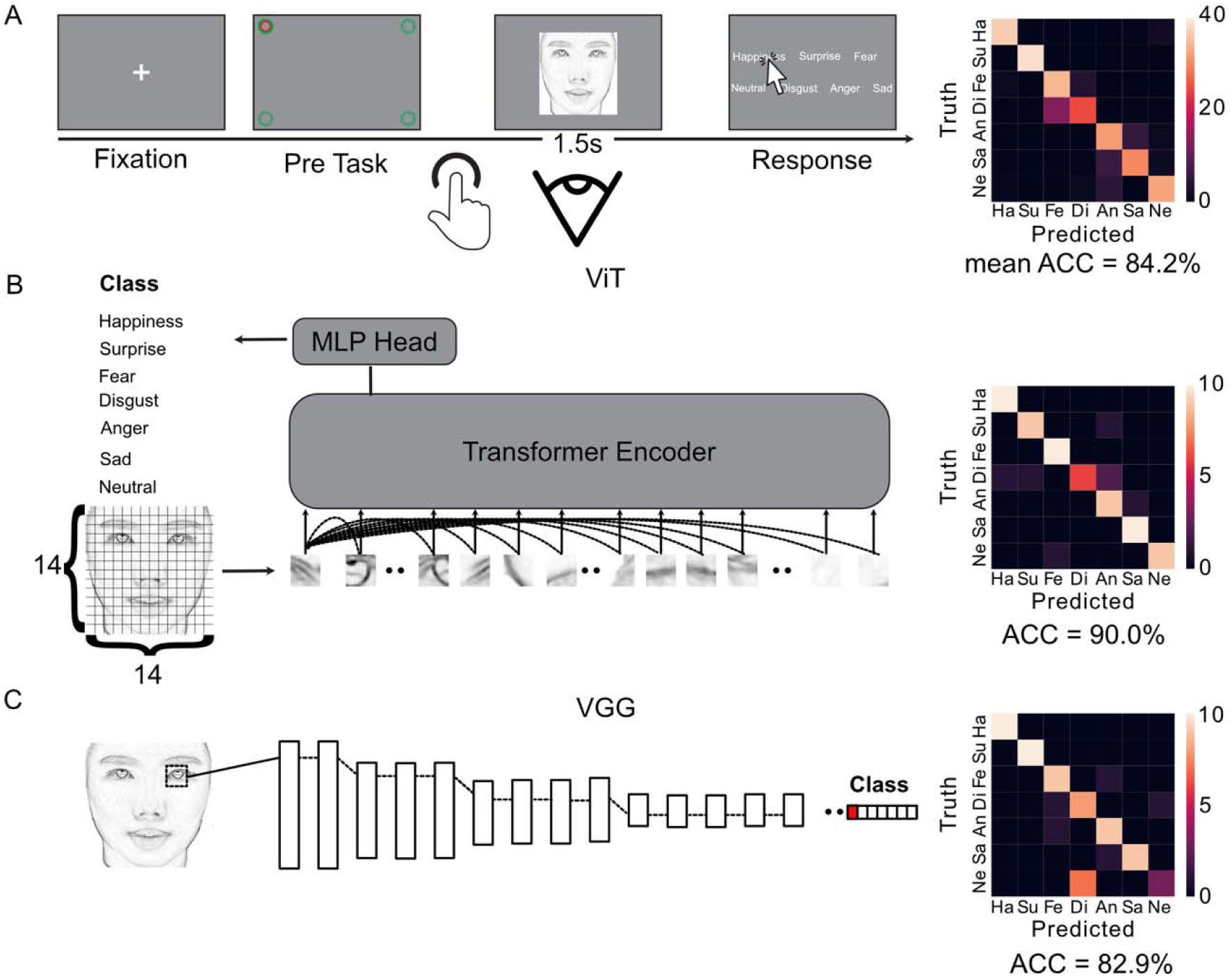
Experiment design and model architectures. (A) Overview of the human experimental procedure for facial expression recognition and the corresponding recognition performance. See the previous study^15^ and the Methods section for details. (B) Architecture of the Vision Transformer (ViT), and its performance in recognizing the same set of facial expressions. (C) Architecture of the Visual Geometry Group (VGG) model and its performance in recognizing the same set of facial expressions.

The two ANNs, VGG and ViT, were first trained and fine-tuned on the 7 emotion categories and then tested on the same set of images used in the human experiment. The images in the test set were not used in the training set. The mean accuracy of VGG was 82.9% and the mean accuracy of ViT was 90.0% (see Figure 1B and 1C for the confusion matrices). While VGG’s prediction performance did not differ significantly from human performance, *t*(27) = 1.22, *p* = 0.234, ViT’s prediction performance exceeded human performance, *t*(27) = 5.38, *p* < 0.001.

### Fixation-related perturbation on emotion recognition performance of ANNs

To investigate if and how fixation-related facial information impacts ANN performance, we applied perturbations to the face images presented to the models. Given that the first two fixations (Fix I and Fix II) play distinct roles in sampling information for emotion recognition^15^, we hypothesized that perturbing facial regions corresponding to Fix I and fix II would differently affect model accuracy. We tested this using two complementary approaches: **information occlusion**, which assessed how prediction accuracy declined as a function of information loss (Figure 2B, upper row), and **information presentation**, which assessed how accuracy improved as a function of added information (Figure 2B, lower row).

**Figure 2.**
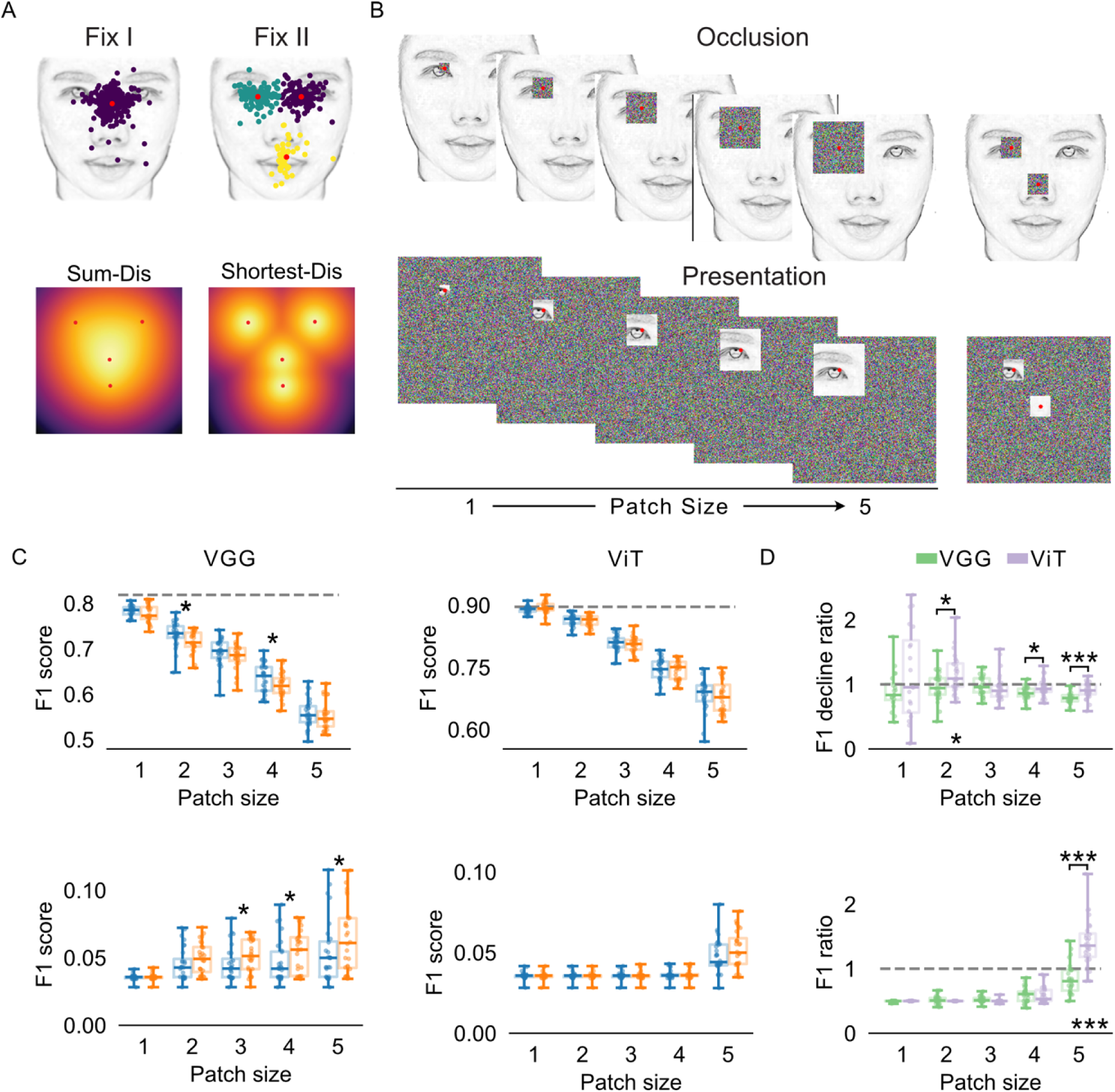
Fixation-related perturbation effects on the ANNs. (A) Distinct roles of Fix I and Fix II in humans. Scatter plots show the clustered distribution of Fix I and Fix II. Heatmaps display the geographical distribution of the summed and shortest distances from each pixel to regions of interest (ROIs), simulating the spatial patterns of the two fixations. (B) Illustration for the patch occlusion and presentations procedures at varying patch sizes. These manipulations were applied to single fixation (left) and paired fixations (right). (C) Results of the fixation-related patch occlusion and patch presentation paradigms applied to the VGG and ViT models. Asterisks indicate significant differences between Fix I-and Fix II-related perturbation effect (* *p* < 0.05) (D) Synergistic effects of paired fixations in patch occlusion and patch presentation on model performance. Asterisks above the paired bars indicate significant differences between Fix I– and Fix II-related synergistic effects, while asterisks above the x-axis indicate significance relative to the baseline – d_R_ / *F_R_* = 1 (* *p* < 0.05, *** *p* < 0.001).

#### Occlusion analysis

For the occlusion analysis, we localized each fixation point on the face image and removed the visual information within a square patch centered at that point. To avoid an arbitrary amount of perturbed information, the patch size was varied from 1 to 5 units of ViT’s standard patch size (16 pixels per patch) and filled with random visual noises. For each of the two ANNs, Fix I-related and Fix II-related occlusions were applied separately to each image. We then calculated the overall *F_1_* scores – a balanced measure of precision and recall – for the occluded images.

In VGG, the prediction accuracy was significantly lower when Fix II-related information was occluded compared to Fix I-related information for patch sizes of 2 and 4 units: 2-unit, *t*(27) = 2.64, adjusted *p* = 0.039; 4-unit, *t*(27) = 2.58, adjusted *p* = 0.039 (FDR-corrected for multiple comparisons; see Figure 2C, upper left). Differences at other patch sizes were not significant (adjusted *p* > 0.23). In ViT, no significant differences were found between Fix I and Fix II occlusions at any patch size (adjusted *p* > 0.44; Figure 2C, upper right).

We next compared the robustness of the two ANNs to occlusion. For each fixation, we calculated performance decline as the ***F_1_* decrease**, defined by subtracting the *F_1_* score of the occluded images from that of the original images. ViT exhibited significantly smaller *F_1_* decreases than VGG for both Fix I and Fix II across all patch sizes (all adjusted *p* < 0.001 after FDR correction), with the smallest *t*(27) = 4.82. These results suggest that ViT is more robust to information loss than VGG.

#### Presentation analysis

For the presentation analysis, we preserved visual information within in a square patch centered at the fixation location, while replacing all other image regions with random visual noise. In VGG (Figure 2C, lower left), the prediction accuracy under Fix II presentation was significantly higher than under Fix I when the patch size exceeded 2 units: 3-unit, *t*(27) = 2.70, adjusted *p* = 0.029; 4-unit, *t*(27) = 3.00, adjusted *p* = 0.028; 5-unit, *t*(27) = 2.30, adjusted *p* = 0.049 (FDR corrected). By contrast, in ViT, the differences between the two fixations were not significant across all patch sizes (all adjusted *p* > 0.36; Figure 2C, lower right).

We also examined which model benefited more from information presentation by comparing *F_1_* scores between the two ANNs. For both fixations, given the same amount of information, VGG outperformed ViT when patch size exceeded 2 units (3 to 5 units). For Fix I, the smallest *t*(27) = 2.08, with the largest adjusted *p* = 0.047 (under 5-unit presentation). For Fix II, the smallest *t*(27) = 3.09, with the largest adjusted *p* = 0.006 (under 5-unit presentation; FDR corrected).

#### Synergistic effects of Fix I and Fix II information

In a further step, we asked if Fix I-related and Fix II-related information jointly produced a synergistic effect on the prediction performance of the two ANNs. To test this, we compared the perturbation effect of the paired information (Fix I + Fix II) with the summed perturbation effects of Fix I and Fix II alone. For the occlusion analysis, we calculated the *F_1_* score decrease for paired occlusion (*d_pair_*), for Fix I alone (*d_I_*), and for Fix II alone (*d_II_*). The synergistic effect was quantified as:

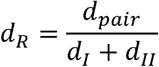

where ***d_R_* = 1** indicates that paired occlusion has the same effect as the sum of the two individual occlusions, and ***d_R_*>** 1 indicates a synergistic effect. As shown in Figure 2D (upper), a *d_R_* > 1 was observed only in ViT with a 2-unit patch size, *t*(27) = 2.29, adjusted *p* = 0.017 (FDR corrected). No such effect was observed in VGG (all adjusted *p* > 0.512). Additionally, ViT showed significantly higher dR values than VGG for patch sizes of 2 units, *t*(27) = 2.58, *p* = 0.026; 4 units, *t*(27) = 2.75, *p* = 0.026; and 5 units, *t*(27) = 5.05, *p* < 0.001 (FDR corrected).

For the presentation analysis, we quantified the synergistic effect using:

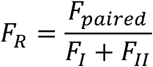

where *F_paired_* denotes the *F_1_* score when both Fix I and Fix II information were presented, and *F_I_* or *F_II_* refers to the *F_1_* score when only Fix I or Fix II information was presented. As shown in Figure 2D (lower), an *F_R_* > 1 was observed only in ViT with a 5-unit patch size, *t*(27) = 5.74, adjusted *p* < 0.001 (FDR corrected). In contrast, VGG showed no such effect (all *t* < 1). Furthermore, under the same 5-unit condition, ViT showed significantly higher FR than VGG, *t*(27) = 7.75, adjusted *p* < 0.001 (FDR corrected).

### The alignment between layer-wise model representations and human fixations

The perturbation results above suggest that ANNs and human eye movements rely on common facial information to perform emotion recognition. To further support this finding, we conducted a layer-wise analysis to examine if – and how – the representations of face images across model layers align with human fixation patterns.

For VGG, we applied Gradient-weighted Class Activation Mapping (Grad-CAN)^25^ to each convolutional layer using the ground-truth facial expression label, generating importance maps for each layer across all images. As shown in the heatmaps in Figure 3A, earlier layers focused on high-frequency features, such as the eyes, mouth, and face outline. In contrast, deeper layers showed more spatially expanded representations of adjacent facial ROIs. Pairwise similarities between layers are shown in the right panel of Figure 3A.

**Figure 3.**
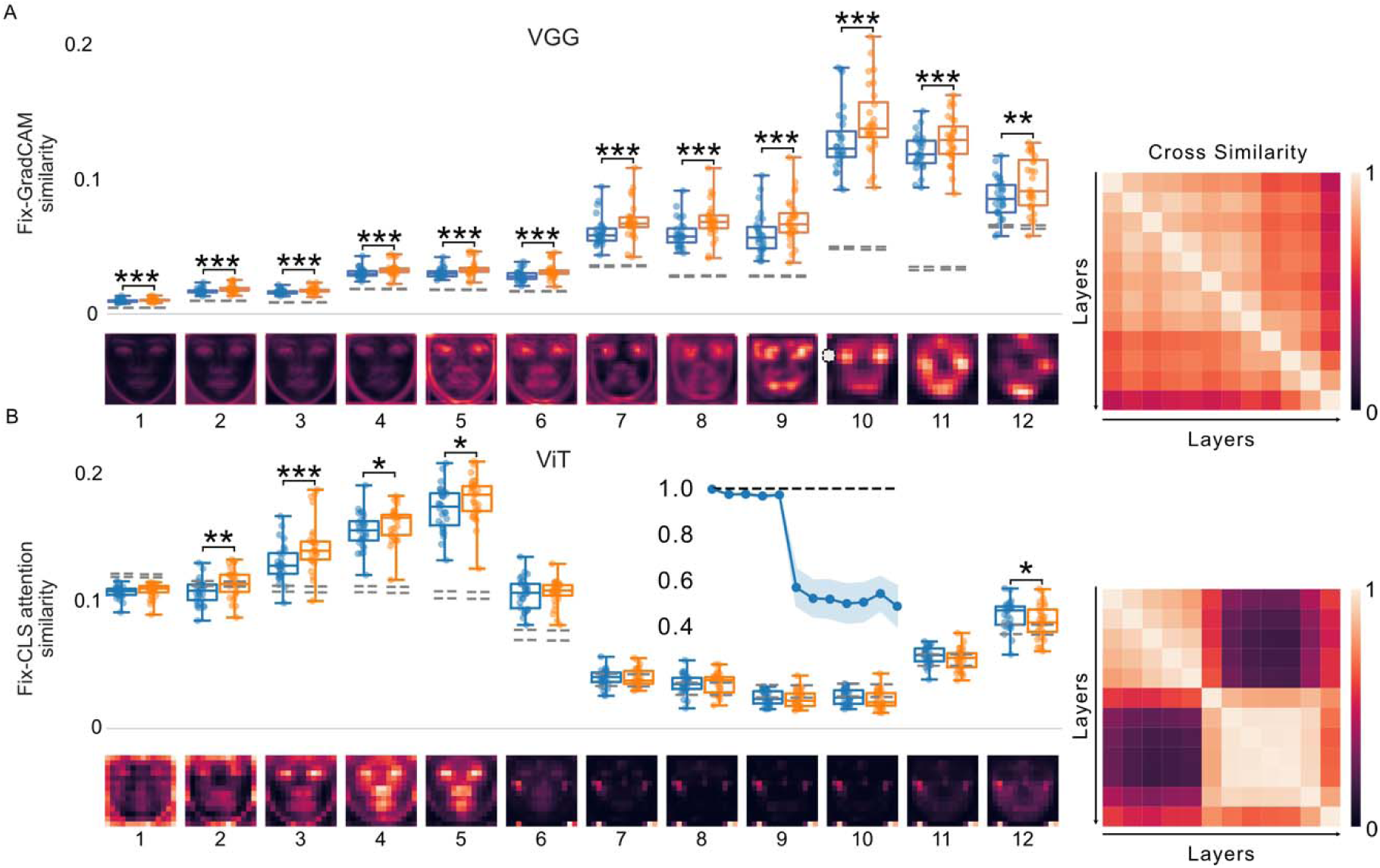
Layer-level correspondence between model importance maps and human fixation distributions. (A) Results of the VGG model, including layer-level importance maps, a between-layer similarity matrix and fixation alignment analysis. (B) Results for the ViT model, featuring attention maps across layers, a similarity matrix, and a line plot showing the representational similarity between the fine-tuned and pre-trained ViT models. Grey dashed lines below the bar plots indicate the upper and lower thresholds of the statistical significance (97.5% and 2.5% of the chance-level similarities distribution, Bonferroni corrected for multiple layers and both fixations). Asterisks indicate significant differences between Fix I and Fix II (\**p* < 0.001, \*\**p* < 0.01, \**p* < 0.05).

To assess VGG’s alignment with human fixations, we generated density maps for Fix I and Fix II and computed the cosine similarity between each fixation map and the corresponding layer’s importance map (boxplots in Figure 3A). For both fixations, all layers showed significant above-chance alignment, with the smallest *t*(27) = 3.61 and the largest adjusted *p* = 0.001 (Bonferroni thresholds were determined by permutation testing; FDR correction was applied for multiple comparisons). Importantly, the importance maps aligned significantly more with Fix II than Fix I, *p*s < 0.001 for all layers after FDR correction, except *t*(27) = 2.84, adjusted *p* = 0.0085 for the final layer. These results are consistent with the perturbation findings, further suggesting that VGG’s information usage is more similar to Fix II than Fix I.

For ViT, we used attention maps as layer-wise representations. In each layer, the attention matrix connected the 1 class token and 196 image patch tokens, indicating how each token attended to all others. We extracted the class token’s attention to the 196 image patches to create a **classification attention map**, which highlighted the regions of the image that contributed most to model predictions. Figure 3B (bottom panel) shows the attention maps averaged across all images. Unlike VGG, ViT’s attention maps were initially diffuse, broadly covering key facial regions. As the layers deepened, attention gradually shifted to more specific patches, culminating in a holistic face shape in the final layer. Pairwise similarities between layers are shown in the right panel of Figure 3B.

To examine ViT’s alignment with human fixations, we calculated the similarity between the fixation density maps and the classification attention maps. For both Fix I and Fix II, layers 3 through 6 showed significant above-chance alignment (all adjusted *p* < 0.0001 after FDR correction), with the smallest *t*(27) = 8.58. As shown in Figure 3B, layers 2 through 5 exhibited significantly higher alignment with Fix II than with Fix I, with *t*(27) values of 3.97, 5.81, 2.63 and 3.24, and adjusted *p* values of 0.003, 0.0004, 0.036, and 0.012, respectively. In contrast, the final layer aligned more closely with Fix I than Fix II, *t*(27) = 2.60, adjusted *p* = 0.036. No significant differences between fixations were observed in the remaining layers.

These results suggest that early layers of ViT attended to task-specific information sampled by Fix II, whereas the final layer attended to more general information associated with Fix I. Notably, ViT’s shift in attention from specific to general information was accompanied by a drop in similarity at the 6^th^ layer (Figure 3B). We speculated that this shift might reflect task-specific representations learned during the pre-training. To test this, we computed the layer-wise representational similarity between the pre-trained and fine-tuned ViT models. As shown in the line plot in Figure 3B, the first five layers already exhibited high similarity between pre-trained and fine-tuned models, followed by a notable drop at the 6^th^ layer.

This pattern suggests that the model’s attention to task-specific information may have originated during pre-training rather than being specific to the model prediction.

### Alignment between representational dependencies in ViT and human fixation transitions

We have so far demonstrated the alignment in information usage between ANNs and individual human fixations. In a third step, we examined the dependencies between sequential fixations. For the human fixation data, we focused on transitions between the first two fixations (Fix I to Fix II) across key facial regions of interest (ROIs): left eye, right eye, nose and mouth (see Figure 4A for ROI definitions and sizes). Fixation III was included as a control to compare Fix I-to-Fix II and Fix II-to-Fix III transitions. Transition probability was defined as the frequency of a fixation in one ROI given a preceding fixation in another ROI.

**Figure 4.**
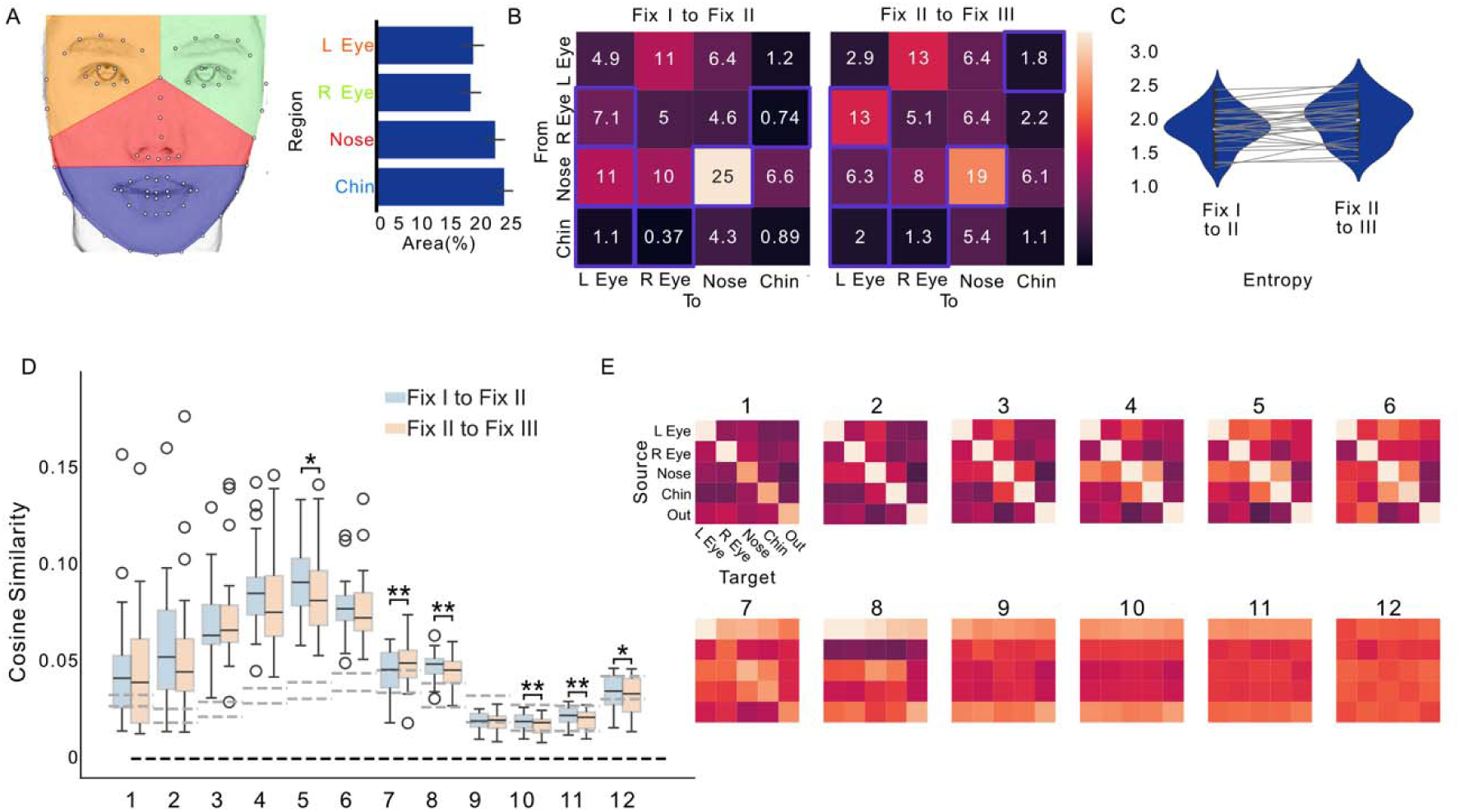
Human fixation transitions and alignment with ViT attention maps. (A) Definition of regions of interest (ROIs) on an example face, with a bar chart showing the average area and standard deviation of each ROI (expressed as a proportion of total ROI area). Color-coded labels correspond to specific facial regions. (B) Group-averaged transition probability matrices for Fix I-to-Fix II and Fix II-to-Fix III transitions. Purple frames highlight ROI pairs with significant differences between the two transition matrices. (C) Entropy values for the two types of transition matrices, with grey lines connecting values of the same participant. (D) Cosine similarity between human fixation transitions (Fix I-to-Fix II and Fix II-to-Fix III) and ViT’s attention maps. Grey dashed lines indicate statistical thresholds (97.5% and 2.5% of the chance similarity distribution, Bonferroni-corrected for multiple layers and transitions). Asterisks indicate a significant difference between the two types of transitions (\**p* < 0.05, \*\**p* < 0.01). (E) Confusion matrices showing average attention weights across the ROIs defined in (A).

As shown in Figure 4B, Fix I-to-Fix II transitions had a significantly higher overall frequency (collapsed across ROIs) than Fix II-to-Fix III transitions, Hotelling’s *T*² test: *t*² = 47.36, *F* = 2.14, *p* = 0.028. Pairwise comparisons revealed significant differences between specific ROI transitions (highlighted by purple frames in Figure 4B), based on t-test with FDR correction. Consistent with our previous findings^15^, Fix I-to-Fix II transitions followed a center-to-divergent pattern, typically moving from the nose to the eyes and mouth. In contrast, Fix II-to-Fix III transitions were more concentrated on non-central ROIs, especially between the eyes.

To quantify transition consistency, we computed the entropy of each transition matrix. As shown in Figure 4C, Fix I-to-Fix II transitions had significantly lower entropy than Fix II-to-Fix III transitions, *t*(27) = 2.78, *p* = 0.0097, indicating a more consistent and structured pattern in the earlier transition. Next, we investigated whether ViT’s attention to inter-patch dependencies aligned with human fixation transitions. For each layer, we computed the cosine similarity between the model’s patch dependency matrix and the human fixation transitions (Fix I-to-Fix II and Fix II-to-Fix III, respectively). As shown in Figure 4D, layers 1-8 exhibited significantly above-chance alignment with both types of transitions (all *p*s < 0.001 after FDR correction), with the smallest *t*(27) = 14.36.

Importantly, layers 5, 7, 8, 10, 11, and 12 showed significant difference in the alignment between Fix I-to-Fix II and Fix II-to-Fix III transitions, most of which had higher alignment with Fix I-to-Fix II transitions. Specifically, layers 5, 8, 10, 11, and 12 showed higher alignment with Fix I-to-Fix II transitions, *t*(27) = 2.55, 3.39, 3.64, 4.25 and 2.65, respectively, *p* = 0.035, 0.007, 0.007, 0.003 and 0.031, whereas only layer 7 showed higher alignment with Fix II-to-Fix III transitions, *t*(27) = 3.35, *p* = 0.007(FDR corrected).

To visualize how the model distributed attention across the image, we grouped attention patches based on the ROIs used in the fixation transition analysis (Figure 4A) and averaged attention weights within each ROI. The resulting confusion matrices (Figure 4E) show that earlier layer exhibited more diffuse attention, with the strongest weights often directed toward the same region (self-directed attention). In contrast, the later layer’s attention was more evenly distributed across different facial regions.

## Discussion

In the present study, we examined the similarity in information usage between the human fixation sequence and two representative ANNs – VGG and ViT – in facial expression recognition. Specifically, we focused on the functional division and collaboration between the initial two fixations in processing facial information and tested if the ANNs exhibited similar properties. We found that both VGG and ViT aligned with the independent function of individual fixations, whereas only ViT showed alignment with the transition and collaboration between fixations. Together, the findings specify the extent to which current ANNs capture the eye movement dynamics involved in facial expression processing and shed light on how the emergence of human-like processing is shaped by the architectural properties of each ANNs.

Our findings showed that both ANNs – VGG and ViT – exhibited alignment in information usage with the two initial fixations, as evidenced by the similarity between fixation patterns and layer-wise representations, as well as by fixation-related perturbation effects on model performance. These result are consistent with prior studies demonstrating representational similarity between ANNs and human face perceptions^7,26,27^. Importantly, the similarity between fixation patterns and ANN representations was consistently higher for Fix II than for Fix I across all convolutional layers of VGG and in the early layers of ViT. Given the functional division between the two fixations – where Fix I processes global, configurational features and Fix II focuses on local, detailed information^15^ – our results suggest a shared mechanism between ANNs and human fixations as feature detectors for face processing. In VGG, all layers were more engaged in processing local detail (Fix II-related), whereas in ViT, early layers showed a similar focus on local features, while the final layer shifted toward global configurational information (Fix I-related).

Beyond the shared alignment with fixation patterns observed in both ANNs, only ViT showed alignment with the transition and collaboration between the two initial fixations. First, ViT – but not VGG – exhibited a synergistic effect when Fix I– and Fix II-related information was perturbed. In both the Information Occlusion and Presentation conditions, perturbing the paired fixation regions had a greater impact on model performance than the summed effects of perturbing each region alone. Second, layers 1-8 of ViT significantly aligned with the transitions between the sequential fixations (e.g., Fix I-to-Fix II and Fix II-to-Fix III). These transitions have been interpreted as a strategy for information processing, in which the preceding fixation guides the location of the subsequent one^19,28^. This statistical dependency between fixation locations mirrors the self-attention mechanism of ViT, which captures dependencies between image patches. Moreover, consistent with prior findings highlighting the importance of the first two fixations over later ones^15,16^, ViT also showed stronger alignment with the Fix I-to-Fix II transition than with the Fix II-to-Fix III transitions in most of the later layers. Third, ViT outperformed VGG in terms of robustness to Information Occlusion and benefit from Information Presentation. This robustness may arise from the self-attention mechanism, which enables ViT to maximize information utility and compensate for missing content through cross-patch interactions. Similar effects have been reported in previous work showing ViT’s minimal performance degradation under partial input loss^29^. Taken together, these findings suggest that ViT aligns with predictive processing and information integration as grounded in the transition and collaboration of human fixation sequences.

The recognition of images – such as faces – during natural visual perception relies on a combination of feedforward and feedback processing in the brain. Both processes are reflected in the fixation patterns observed during image viewing. The feedforward processing is driven by the physical features of the visual stimuli^30,31^ and can be modeled as a saliency map that guides the allocation of fixation locations^32^. In contrast, the feedback processing is guided by top-down predictions about incoming visual information^33,34^ and is reflected in the sequences of fixations over time^35,36^. In our results, these two types of processing – feedforward and feedback – were distinctly captured by the two ANN models: VGG, which reflects feedforward processes of the visual ventral pathway, and ViT, whose self-attention mechanism functions as a form of global feedback processing.

Although ViT aligned with the human fixation sequence in representing and integrating task-relevant facial information, the underlying mechanisms may differ between ViT and human fixation sequence. Prior research has shown that the sampling of global, configurational facial information typically precedes the sampling of local fine details^37^, corresponding to Fix I and Fix II, respectively^15^. The coarse-to-fine sequence supports a processing pipeline in which an initial hypothesis is formed about the task-relevant dimension of the face (e.g., identity or emotion), which then guides subsequent information sampling to test the hypothesis and generate a behavioral response^18^. However, in ViT, early layers showed greater alignment with Fix II than Fix I, while the final layer showed the opposite pattern – greater alignment with Fix I than Fix II. Moreover, the graphical patterns of layer-wise representations revealed that both the processing of local details (Figure 3B, bottom panel) and the transition between face ROIs (Figure 4E) occurred predominantly in early layers, but less so in later ones. These results suggest that ViT processes local fine-grained information before integrating general configurational features – an order opposite to that of human eye movements.

This reversed order may reflect how ViT is structured: early layers are optimized for local feature extraction, while later layers for global integration. The discrepancy between ViT and human visual processing raises interesting questions, especially when considering ViT’s superior performance over humans in the emotion recognition task. One possible explanation is that ViT’s fine-to-coarse processing pipeline is optimized for the specific demands of emotion recognition, the specific task it was fine-tuned for. However, this task-specific optimization might come at a cost to performance in other tasks, such as identity recognition, which may rely on a different balance of local and global information. In real-world scenarios, face perception is not limited to a single predefined task goal but often requires flexible switching between task goals. A coarse-to-fine processing strategy, as seen in human vision, may therefore reflect an adaption for balancing task flexibility with specificity. The differences observed here open important questions for future research regarding the relative strengths and limitations of the two-stage pipeline.

A growing body of research has demonstrated alignment between ANNs and human neural processing of visual objects^26,27,38^, as well as the reproduction of many human-like behaviors in face recognition tasks^6–8,39^. While previous studies using convolutional neural networks have treated the entire image as input and hierarchically extracted features layer by layer, it remained unclear what specific features were used and how they contributed to recognition. By comparing ANNs with human fixation sequences, the current findings reveal both shared and distinct mechanisms of information usage across the two models in facial expression recognition. These insights enhance the interpretability of ANNs as models of human cognition and suggest that incorporating human-like fixation strategies may benefit task-specific performance. For example, an ANN trained to predict human duration perception of video clips performed better when the model input was constrained by human fixation locations rather than when the full video content was used^40^. Similarly, to reduce the computational cost the Vision Transformer, human fixation patterns have been used to pre-select patches during training^41^.

One limitation of our study is that the alignment between ANNs and human fixation patterns was examined only in the context of emotion recognition. We did not include identity recognition, as in our previous eye movement study, in order to avoid the risk of model overfitting due to the large number of identity involved (n = 70, 15). As a result, it remains unclear if the current findings can generalize to other face recognition tasks. A second limitation is that our models were trained and evaluated in a single-goal task setting, rather than in a multiple-goal context (e.g., recognizing both identity and emotion). This may limit the extent to which the ANNs align with human face processing, which in real-life scenarios often involves concurrent or dynamically shifting task goals.

## Methods

### Experiment Design

Eye movement data were curated from our previous study^15^, in which participants completed both an emotion recognition task and an identity recognition task in a counterbalanced order. To avoid model overfitting due to the large number of identities (n = 70), only the data from the emotion recognition task were analyzed in the present study.

#### Participants

Twenty-eight healthy college students (18 females; mean age = 23.2 years, SD = 2.3) participated in the experiment. All participants had normal or corrected-to-normal vision. The study was conducted in accordance with the Declaration of Helsinki and was approved by the Ethics Committee of the School of Psychological and Cognitive Sciences, Peking University (Approval #2020-10-01).

#### Stimuli and procedure

The stimuli consisted of 70 front-view facial images selected from the Chinese Facial Affective Picture System^42^, with 10 images representing each of the 7 emotion categories. On each trial, participants freely viewed a face image for 1500 ms, during which eye movements were recorded. Participants were asked to choose one of seven emotion labels to match the perceived emotion of the image. For further details, see^15^.

#### Analysis of behavioral Data

For each participant, a 7×7 confusion matrix was computed to quantify emotion recognition accuracy. Each row represented the ground truth (i.e., the actual emotion category), and each column represented the participant’s chosen label. The group-level confusion matrix was obtained by averaging individual matrices across all participants.

#### Region-of-interest (ROI) Definition

Facial landmarks for each image were detected using the machine-learning toolkit Dlib with a pretrained model^43^. Based on these landmarks, four ROIs—left eye, right eye, nose, and mouth—were defined with constraints to balance the area of each ROI^44,45^. The size of each ROI was quantified using the Shapely package^46^.

#### Fixation transition across ROIs

For each participant, a 4×4 transition matrix was computed to capture the probability of transitions between consecutive fixations across the four ROIs. Rows in the matrix represented the ROI of the preceding fixation, while columns represented the ROI of the subsequent fixation. Each cell contained the probability of the next fixation landing in a specific ROI, given the location of the preceding fixation. Group-level transition matrices were obtained by averaging individual matrices across all participants. Transitions were computed separately for Fix I to Fix II and Fix II to Fix III.

#### Entropy of Fixation Transition

To quantify the consistency of fixation transition patterns, we calculated the Shannon entropy^47^ of each participant’s transition matrix. Specifically, each 4×4 matrix was flattened into a one-dimensional vector, and entropy was computed using the formula:

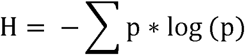

where p represents the transition probability in each cell of the matrix. Entropy was calculated separately for Fix I-to-Fix II and Fix II-to-Fix III transitions. A paired t-test was used to compare the entropy values between the two types of transitions across participants.

### Artificial neural networks (ANN) models

#### VGG model architecture

The VGG16 model used in this study consisted of 13 convolutional layers followed by 3 fully connected layers. Each convolutional layer employed 3×3 filters and was followed by max-pooling layers to reduce the spatial dimensions of the feature maps. The model was pre-trained on the ImageNet 2012 dataset^48^, which contains over 1 million images across 1,000 object categories. To adapt the model for facial emotion recognition, we replaced the original 1,000-class output layer with a new fully connected layer designed to classify the 7 emotion categories. The adapted model was then fine-tuned on our emotion-labeled dataset. The first convolutional layer (Layer 0) was excluded from subsequent analyses, as it primarily encodes low-level visual features.

#### ViT model architecture

The Vision Transformer (ViT) model follows a transformer-based architecture. For each input image, ViT divides the image into fixed-size patches, treating them as a sequence of tokens. The model computes the statistical relationships among these patches using self-attention mechanisms, enabling a global understanding of the visual input. In this study, we used the **vit-base-patch16-224** architecture developed by Google. This model includes 12 transformer layers, each with 12 attention heads and a hidden size of 768, meaning each patch is represented by a 768-dimensional feature vector. Similar to the VGG model, ViT was pre-trained on the ImageNet 2012 dataset^48^, which includes 1 million images across 1,000 categories. To adapt the model for our emotion recognition task, we replaced the final classification layer with a new fully connected layer for the 7 emotion categories, followed by fine-tuning on our training dataset.

#### Model fine-tuing and prediction

To fine-tune both the VGG and ViT models, we constructed a combined training dataset comprising images from the FER+ dataset^49^, the Extended Cohn-Kanade dataset(CK+)^50^, and the other images from the Chinese Facial Affective Picture System^42^ that were not included in the human experiment. All images in this dataset were labeled with one of the seven emotion categories used in our study.

To ensure consistency with the emotion categories in the human experiment, we excluded images labeled as “contempt” from FER+. A validation set was constructed by combining the official validation set from FER+ and 20% of the images from CK+.

Both VGG and ViT were fine-tuned on this combined dataset using the adapted architecture described above. After fine-tuning, each model was tested on the exact set of images presented to human participants in the experiment. For each model, a 7×7 confusion matrix was computed to evaluate prediction performance across emotion categories.

### Statistical information

#### Perturbation Analysis

We conducted two types of perturbation analyses—Occlusion and Presentation—to evaluate how fixation-related facial information influenced model performance. The Occlusion condition assessed how removing visual information affected prediction accuracy, whereas the Presentation condition assessed how adding isolated visual information influenced accuracy.

To match the structure of the ViT model, each image was divided into a 14×14 grid of patches, with each patch measuring 16×16 pixels. Fixation-related patches were determined by matching each human fixation point to the nearest patch center. Perturbation was applied to a variable number of patches around each fixation point, ranging from 1 to 5 contiguous units, allowing us to simulate foveated sampling regions of increasing size. Based on the image dimensions and experimental setup, one patch unit corresponded to approximately 0.5° of visual angle, meaning that 5 patch units approximated 2.5°, which falls within the typical range of the eye’s foveal high-acuity region^51,52^.

In the **Occlusion** condition, the target patch region was masked by random visual noise (pixel intensity: mean = 128, SD = 64 in 8-bit scale), simulating information loss. Prediction accuracy was then measured for each model on the occluded images.

In the **Presentation** condition, only the patch region corresponded with the fixation was preserved, and the remaining image content was replaced with the same random visual noise. Prediction accuracy was measured to evaluate how well the isolated region supported classification.

#### Layer-wise analysis of model representations

For each convolutional layer in the VGG model, we applied Gradient-weighted Class Activation Mapping (Grad-CAM)^25^ to generate importance maps for face classification. These maps reflected the contribution of spatial features in each layer toward predicting the ground-truth emotion label. Activation maps were computed for each face image used in the human experiment.

In the ViT model, each transformer layer produced a self-attention map, which quantified the contribution of image patches to classification through attention weights centered on the [CLS] token. We extracted the attention from [CLS] to each of the 196 image patches as the layer’s classification attention map, which was treated as the activation map for analysis.

#### Alignment with Human Fixation Maps

To assess the alignment between model representations and human eye movements, we computed the cosine similarity between each model’s activation map and the fixation density maps derived from human Fix I and Fix II, separately for each image. For each layer, this analysis was conducted across all images and participants.

To evaluate statistical significance, we implemented a two-step permutation procedure:

1. Within-participant permutations: For each participant, we shuffled the activation maps and computed their cosine similarity with the fixation density maps, repeating the process 1,000 times to generate a null distribution of similarity values.
2. Group-level sampling: These individual-level null distributions were pooled across participants and resampled 10,000 times to generate group-level null distributions.

From the null distribution, we derived Bonferroni-corrected thresholds for significance (α = 0.05 / 24, accounting for 12 layers × 2 fixations). Independent t-tests were performed to compare observed cosine similarities with chance thresholds, followed by FDR correction for multiple comparisons. We also conducted paired t-tests between Fix I and Fix II similarity scores at each layer to identify fixation-specific alignment.

#### Inter-Patch Dependency and Fixation Transition Alignment

To assess if ViT’s attention mechanism mirrored human fixation transitions, we extracted inter-patch attention matrices from each layer. For each image and layer, a 12×197×197 matrix was obtained, where 197 tokens represented [CLS] + 196 patches, and 12 attention heads reflected diverse attention patterns. We averaged attention weights across heads to generate a 196×196 dependency matrix indicating the strength of dependence between each pair of patches. To match the directionality of fixation transitions (e.g., Fix I → Fix II), the matrix was transposed.

We then computed the cosine similarity between each inter-patch dependency matrix and the fixation transition matrices for Fix I-to-Fix II and Fix II-to-Fix III. The same two-step permutation procedure described above was used to assess the statistical significance of alignment, with separate evaluations conducted for each layer and each transition.

## Acknowledgements

This work was supported by the National Natural Science Foundation of China (Grant No. 32271086) awarded to LW and ML. Additional support was provided by the Mercator Fellowship of the Deutsche Forschungsgemeinschaft (Grant No. 450600965) to LW and by the Shanghai Jiao Tong University Grant (Grant No. YG2025QNB13) to LW.

